# Spatial control of mitochondrial retrograde signaling by nuclear pore-associated contact sites

**DOI:** 10.64898/2026.07.23.740218

**Authors:** Sahana Mitra, Giulia Burrone, Surya Rubarajan, Christophe Klein, Tyler Kwok, Suchit Gutta, Keyu Chloe Li, Elizaveta Berson, Joseph Orofino, Maria Dafne Cardamone, Michael Blower, Valentina Perissi

## Abstract

Mitochondria relay their functional state to the nucleus via retrograde signaling, yet whether the spatial organization of the mitochondrial network plays a role in this process remains unclear. Here, we show that stress-induced clustering of mitochondria around the nucleus is a crucial part of the retrograde response. Perinuclear clustering facilitates the formation of mitochondria– nucleus contact sites (MNCS) and the nuclear entry of GPS2, a key mediator of mitochondrial retrograde signaling essential for activating nuclear-encoded mitochondrial and stress-response genes in response to various mitochondrial stressors. Unexpectedly, TSPO-driven MNCS are dispensable for GPS2-based retrograde signaling. Instead, we identify the mitochondrial import receptor TOMM70 and the nucleoporin RanBP2/NUP358 as components of a stress-induced nuclear pore-associated tethering complex required for promoting GPS2 nuclear translocation and activation of downstream programs. These findings establish MNCS as a functional gateway for mitochondrial retrograde signaling, highlighting that organelle positioning and tethering at the nuclear pore provide an unexpected layer of stress regulation.

## Introduction

Mitochondria are versatile organelles essential for maintaining cellular homeostasis, energy production, and adapting to environmental stresses. Although they contain their own DNA, most mitochondrial proteins are encoded by nuclear genes and imported into the organelle after translation^1–3^. This evolutionary reliance emphasizes the importance of effective bidirectional communication: anterograde signals from the nucleus regulate mitochondrial biogenesis and function, while retrograde signals from mitochondria inform the nucleus of their status, triggering adaptive gene expression^1,4^. Mitochondrial dysfunction caused by genetic mutations, metabolic challenges, or environmental toxins activates a mitochondrial stress response (MSR), also known as the mitochondrial integrated stress response (ISRmt) or the mitochondrial unfolded protein response (UPRmt), to reprogram nuclear gene activity as needed to maintain cellular homeostasis^5,6^. Malfunctions in these communication networks contribute to mitochondrial disorders, metabolic diseases, neurodegeneration, and aging, highlighting the critical role of coordinating the two genomes within a cell^7,8^.

Mitochondria communicate their functional status to the nucleus through a range of retrograde messengers and signaling pathways. Traditional signals include shifts in metabolite and second messenger levels, such as reactive oxygen species (ROS), calcium, and the AMP/ATP ratio, which together modulate the activity of nuclear transcription factors and chromatin-modifying enzymes^4,7^. More recently, the DELE1 was characterized as a direct relay linking mitochondrial membrane depolarization and protein import stress to phosphorylation of eIF2alpha and ATF4-driven gene transcription^9–13^. We also identified G-Protein Pathway Suppressor 2 (GPS2) as a direct mediator of mammalian retrograde signaling in response to mitochondrial depolarization^14^. GPS2, a multifunctional transcriptional cofactor present in both mitochondria and the nucleus^15–20^, translocates from mitochondria to the nucleus to promote the activation of mitochondrial and stress response genes by facilitating H3K9 demethylation and RNA Polymerase II activity through inhibition of Ubc13-mediated K63-ubiquitination^14,21,22^. GPS2-mediated retrograde signaling is essential for adaptive gene expression in response to mitochondrial stress, for promoting mitochondrial biogenesis in differentiating adipocytes and brown adipose tissue, and for promoting fatty acid oxidation in liver stellate cells^14,23^, thus confirming its role as a key mitochondrial-to-nucleus communication pathway in mammals.

Despite significant advances in understanding retrograde signaling at the molecular level, its spatial regulation within cells is still not well understood. Mitochondria are highly dynamic, mobile organelles. During stress, they tend to cluster around the nucleus, a process known as perinuclear clustering. Although stress-induced perinuclear enrichment of mitochondria has been documented across diverse cell models, its role in retrograde communication is unclear. Key questions remain: is physical proximity between mitochondria and the nucleus required for signaling? And is the positioning of the mitochondrial network a controlled aspect of the retrograde response, or just a consequence of dysfunction? This distinction has broad implications: if proximity is crucial, then mechanisms that govern mitochondrial placement become essential components of the stress response.

Membrane contact sites (MCS) act as essential hubs for communication between organelles in eukaryotic cells. These regions, where membranes are closely aligned, usually 10-80 nm apart, are stabilized by tethering proteins. Short-range contact sites offer more efficient, specific, and controllable platforms for signaling than long-distance diffusion. They enable functions such as lipid transfer, Ca2+ exchange, organelle movement, and signal transduction, and their disruption can lead to disease ^2,24,25^. Mitochondria-associated membranes (MAMs), or mitochondria-ER contact sites (MERCs), are best known for regulating phospholipid biosynthesis, Ca2+ signaling, and mitochondrial fission at the interface between mitochondria and the ER^26–29^. In addition to the ER, mitochondria form regulated, functional contacts with lysosomes, lipid droplets, peroxisomes, and the plasma membrane^30–33^. Evidence of physical interactions between mitochondria and the nuclear envelope has accumulated over decades^34,35^, but their functional importance is only now being explored. The development of Split-GFP-based Contact Site sensors (SPLICS) has enabled the measurement and analysis of mitochondria-nucleus contact sites (MNCS) in live cells^36–38^, providing an opportunity to screen for and assess the role of putative mitonuclear tethering factors^39,40^. In yeast, a high-throughput colocalization screen identified Cnm1 as the primary nuclear tether for MNCS, working alongside the outer mitochondrial membrane transporter Tom70^41^. In mammalian cells, Mitofusin-2 (MFN2), TSPO, ACBD3, and AKAP95 have been linked to MNCS formation^42–44^. Most importantly, these studies begin to imply an active role for MNCS in promoting mitonuclear communication via NF-kB signaling^39,42,44^ and, more recently, in supporting the nucleus’s energetic needs via import of ATP and phosphocreatine^45^. Nonetheless, it remains unaddressed whether MNCS are broadly and functionally required for mitochondrial retrograde signaling, particularly for the direct transfer of protein cargo from mitochondria to the nucleus in response to mitochondrial stress.

Here, we demonstrate that stress-induced perinuclear clustering promotes the formation of mitochondria–nucleus contact sites (MNCS), which support GPS2-dependent retrograde signaling. Our results suggest a model where the mitochondrial import receptor TOMM70 and the nucleoporin RanBP2 work together at a nuclear pore-associated contact site, orchestrating stress-induced communication from mitochondria to the nucleus.

## Results

Perinuclear clustering is a well-known phenomenon observed in response to stress and matrix stiffness, as well as during epithelial-mesenchymal transition^46–49^. However, its functional relevance remains unclear: some studies suggest it promotes mitonuclear communication, while others consider it a side effect of dysfunctional mitochondria that collapse onto the nucleus and fail to provide energy to peripheral locations. To formally address this question, we evaluated the extent to which perinuclear clustering is required for mitochondrial retrograde signaling. As a readout, we used GPS2 nuclear accumulation, which we previously showed depends on SUMO-regulated retrograde translocation from uncoupled mitochondria to the nucleus^14^. Because our previous studies were limited to GPS2 translocation in response to FCCP-induced depolarization in 3T3-L1 cells, we first investigated whether the same increase in nuclear accumulation could be recorded in response to other stressors and across distinct cell types.

Our results confirm that in 3T3-L1 cells, significant nuclear accumulation of GPS2 was observed within 3 h of treatment not only with FCCP but also with the Complex III inhibitor Antimycin (**Figure 1A)**. An even more robust response was observed when mitochondrial protein import was directly inhibited with MitoBloCK-6 (MB6)(**Figure 1B**), suggesting that depolarization-induced retrograde transport is likely favored by reduced protein import efficiency into mitochondria as previously reported for other retrograde factors^50^. GPS2 nuclear accumulation in response to the Complex I inhibitor Rotenone was also slightly increased at 3 h, although not significantly (**Supp Figure S1A)**. Significant nuclear accumulation of GPS2 was also observed in human U2OS cells in response to 1 h treatment with FCCP, MB6, and the ATP synthase inhibitor Oligomycin (**Figure 1C**). Notably, in the case of Rotenone, significant nuclear accumulation was not observed until 2 h after treatment (**Supp Figure S1B**), suggesting that, in both human and mouse cells, the kinetics of GPS2 signaling activation in response to Complex I inhibition are slower than those for other mitochondrial toxins. Together, these results indicate that GPS2 retrograde signaling is activated in response to multiple sources of mitochondrial stress, with only minimal differences in the timing of accumulation likely dependent on the impact of different stressors on membrane potential kinetics and depolarization-induced inhibition of protein import into the mitochondria, thereby supporting its relevance as a measurable outcome of mitonuclear communication across human and murine cell models.

**Figure 1.**
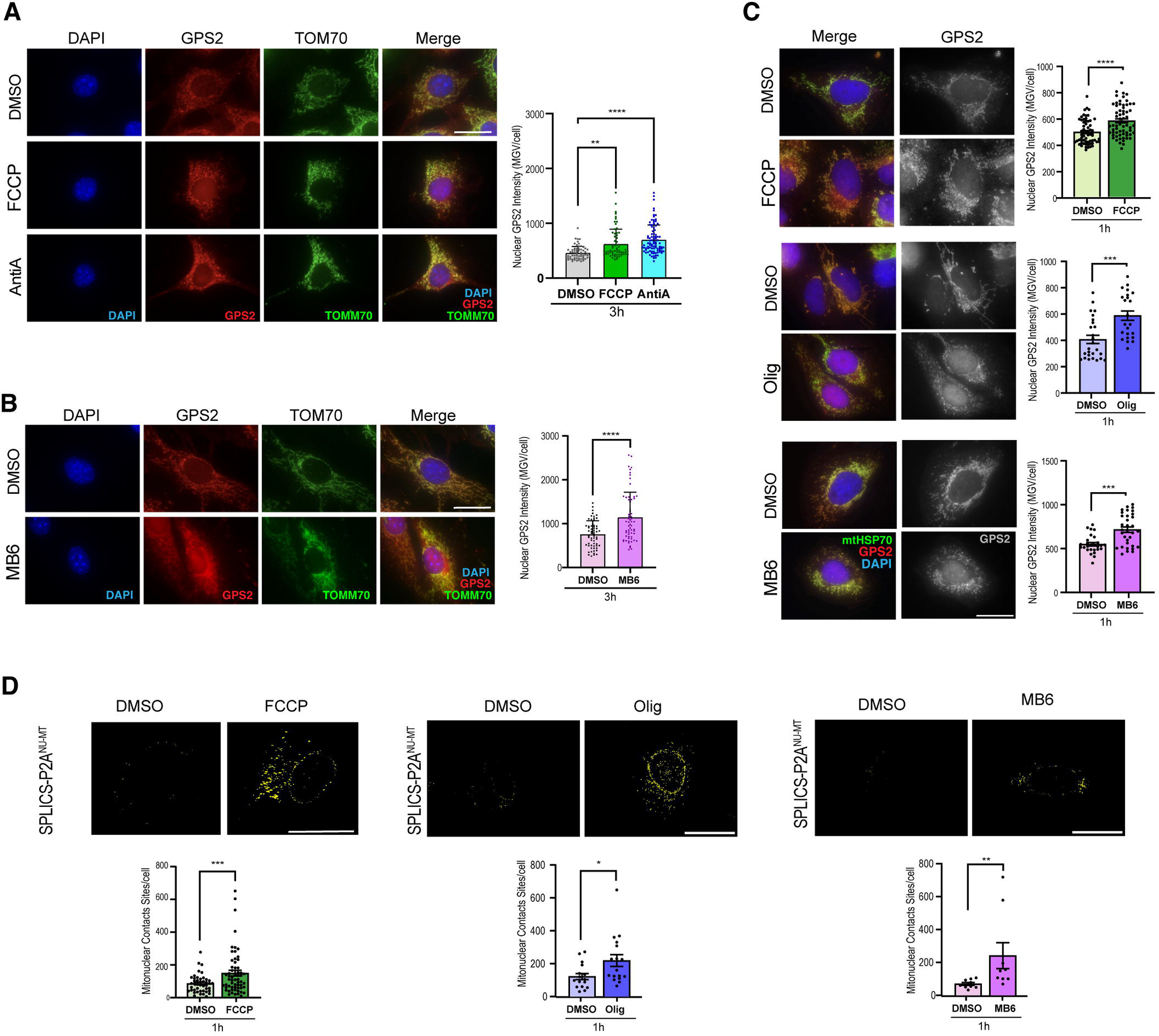
Diverse mitochondrial stressors drive GPS2 nuclear accumulation, perinuclear clustering, and formation of mitochondria–nucleus contact sites (MNCS). (A) Representative immunofluorescence images of 3T3-L1 cells treated for 3 hours with FCCP 10µM, Antimycin A (AntiA) 10µM, or vehicle control (DMSO) stained for DAPI (blue), GPS2 (red), and TOMM70 (green); merge and individual channels shown. Right, quantification of nuclear GPS2 intensity (mean gray value, MGV per nucleus). Statistical analysis of the data was performed using Ordinary one-way ANOVA. Statistical significance was defined as follows: p-value 0.1234 (ns), 0.0332 (*), 0.0021 (**), 0.0002 (***), and <0.0001 (****). The scale bar represents 20 µm. (B) Representative immunofluorescence images of 3T3-L1 cells treated for 3 hours with MitoBloCK-6 (MB6) 50µM or vehicle control (DMSO) stained for DAPI (blue), GPS2 (red), and TOMM70 (green); merge and individual channels shown. Right, quantification of nuclear GPS2 intensity (mean gray value, MGV per nucleus). Statistical analysis of three independent experiments was performed using unpaired t-test. Statistical significance was defined as follows: p-value 0.1234 (ns), 0.0332 (*), 0.0021 (**), 0.0002 (***), and <0.0001 (****). The scale bar represents 20 µm. (C) Representative immunofluorescence images of U2OS cells treated for 1 hour with DMSO, FCCP, Oligomycin (Oligo), or MB6, stained for mtHSP70 (green), GPS2 (red), and DAPI (blue); merge and single-channel GPS2 (gray scale) shown. Right, quantification of nuclear GPS2 intensity (MGV per nucleus) for each stressor versus its paired DMSO control. Data points in the adjacent graph represent individual nuclei. For FCCP treatment, DMSO, *n* = 61; FCCP, *n* = 73; For Oligomycin treatment, DMSO, *n* = 26; Oligo, *n* = 23; For MB6 treatment, DMSO, *n* = 27; MB6, *n* = 33. P values were calculated using the two-tailed *t*-test. Data are presented as mean ± SEM from three independent experiments, * p <0.05, ** p < 0.01, *** p <0.001. The scale bar represents 20 µm. (D) Mitochondria–nuclear contact sites visualized in U2OS cells expressing the split-GFP mito-nuclear reporter SPLICS-P2A^NU-MT^ (reconstituted GFP signal shown as a 3D rendering of the complete optical stack, yellow), after 1 hour treatment with DMSO, FCCP, MB6, and Oligomycin. Right, quantification of MNCS per cell. Each dot represents one analyzed cell, identified by its corresponding nuclear ROI. Only cells with adequate SPLICS reporter expression and image quality for reliable three-dimensional rendering and contact-site quantification were included. For FCCP treatment, DMSO, *n* = 54; FCCP, *n* = 63; For Oligomycin treatment: DMSO n= 16, Olig= 17; For MB6 treatment, DMSO n=9, MB6 n=9. P values were calculated using a two-tailed *t*-test. Data are presented as mean ± SEM from three independent experiments, *<0.05, **< 0.01 and ***<0.001. The scale bar represents 20 µm.

As expected, under all tested stress conditions and across both cell models, we observed mitochondrial fragmentation and perinuclear clustering of fragmented mitochondria (**Figure 1A-1C**). To assess whether the spatial reorganization of stressed mitochondria to the perinuclear space promotes the formation of mitonuclear contact sites (MNCS), we employed a mitonuclear Split-GFP-based Contact Site Sensor (SPLICS-P2A^MT-NU^) reporter, previously developed to monitor NF-κB translocation through MNCS in response to TNF-α^39^. Fluorescence signal reconstituted within the perinuclear space when the outer mitochondrial membrane (OMM)-targeted GFP_1-10_ and outer nuclear membrane (ONM)-targeted GFP-β11 are in proximity was acquired and quantified as described before^36^. Results show that a short, 1 h treatment with either FCCP, Oligomycin or MitoBlock6 is sufficient to drive a significant increase in MNCS in U2OS cells (**Figure 1D**), in the absence of any gross morphological change in the nuclear envelope structure (**Supp Figure S1C**). Although the transient expression of the SPLICS-P2A^MT-NU^ reporter shows as expected some minimal leakage to the ER^51^, the increase in GFP reconstituted signal upon mitochondrial stress is specific to mitonuclear contacts, as indicated by i) the fact that the quantification of contact sites/cells is limited to the perinuclear area as previously described^36^ (**Figure 1D**); and ii) is not observed when specifically examining mito-ER contact sites with the SPLICS-P2A^MT-ER^ reporter^37^ (**Supp Figure S1D**).

Once established that mitochondria-stress-driven relocalization of GPS2 to the nucleus correlated with increased perinuclear mitochondrial clustering and the formation of MNCS in response to multiple stressors, we set out to investigate the extent to which proximity was required for signaling. First, we impaired perinuclear clustering by transiently downregulating either BAG6 or p62/SQSTM1. Previous studies had shown a clear role for both proteins in stress-induced perinuclear clustering^48,52^. As expected, downregulation of either protein was sufficient to inhibit FCCP-induced perinuclear clustering in HeLa cells (**Figure 2A**, **2B**). Quantification of GPS2 nuclear accumulation revealed that FCCP-induced GPS2 retrograde translocation following a 1 h treatment was inhibited when clustering was impaired by downregulating either BAG6 or p62 (**Figure 2C, 2D**). As our previous studies showed that nuclear translocation of GPS2 was essential for the transcriptional activation of nuclear-encoded mitochondrial genes and stress response genes^14^, we also probed the impact of BAG6/p62 downregulation on a representative subset of GPS2 direct targets, including *AKAP1, ASNS, LONP1, NDUFV1*. RT-qPCR results confirmed that FCCP-induced activation of each target is blunted when perinuclear clustering is impaired (**Figure 2E**). Conversely, downregulating the motor protein KIF5B, which was previously shown to be essential for maintaining the mitochondrial network extended across the cytosol^53^, led to mitochondria collapsing into the nucleus (**Figure 2F**), retrograde translocation and nuclear accumulation of GPS2 (**Figure 2F),** and activation of GPS2 stress response targets even in the absence of FCCP treatment (**Figure 2G)**. Similar results were observed in both HeLa and U2OS cells (**Figure 2H).**

**Figure 2.**
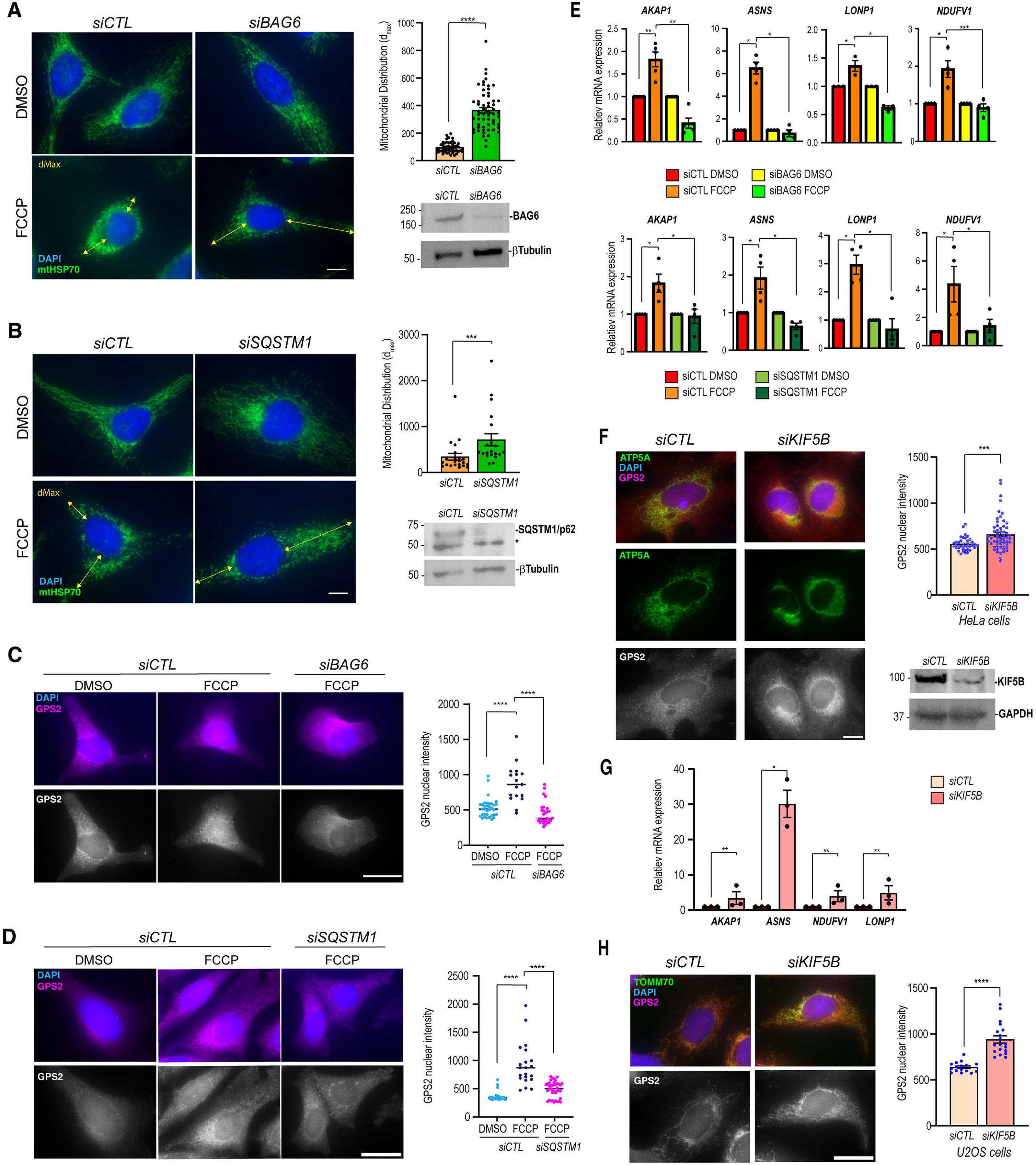
Perinuclear clustering is required for GPS2 retrograde translocation and stress-gene induction, and forced perinuclear collapse is sufficient to trigger signaling. (A) Representative immunofluorescence images of HeLa cells transfected with control (si*CTL*) or BAG6 (si*BAG6*) siRNA and treated with DMSO or FCCP (10 µM for 1 hour), stained for mtHSP70 (green) and DAPI (blue); yellow lines indicate the maximal radial distance of mitochondrial distribution (dMax). The scale bar represents 20 µm. Middle, quantification of mitochondrial distribution (dMax). Data points in the adjacent graph represent individual nuclei. For si*CTL*, *n*= 48; si*BAG6*, *n*= 68. P values were calculated using the two-tailed *t*-test (**** represents p< 0.0001). Data are presented as mean ± SEM from three independent experiments. Right, representative immunoblot for BAG6 with β-tubulin as loading control. (B) As in (A) for HeLa cells transfected with control or p62/SQSTM1 (si*SQSTM1*) siRNA. Middle, quantification of mitochondrial distribution (dMax). The scale bar represents 20 µm. Data points in the adjacent graph represent individual nuclei. For si*CT*L, *n*= 28; si*SQSTM1*, *n*= 28. P values were calculated using the two-tailed *t*-test (*** represents p< 0.001). Data are presented as mean ± SEM from three independent experiments. Right, immunoblot for SQSTM1/p62 (asterisk denotes nonspecific band) with β-tubulin. (C) Representative immunofluorescence images of HeLa cells (si*CTL* + DMSO, si*CTL* + FCCP, si*BAG6* + FCCP) stained for DAPI (blue) and GPS2 (magenta); single-channel GPS2 (gray scale) shown. The scale bar represents 20 µm. Right, quantification of nuclear GPS2 intensity. Data points in the adjacent graph represent individual nuclei. For si*CTL* + DMSO, *n*= 34; si*CTL* + FCCP, *n*= 17; si*BAG6* + FCCP, *n*= 30. P values were calculated using the two-tailed *t*-test (**** represents p< 0.0001). Data are presented as mean ± SEM from three independent experiments. (D) As in (C) for control versus si*SQSTM1*. For si*CTL* + DMSO, *n*= 36; si*CTL* + FCCP, *n*= 21; si*SQSTM1* + FCCP, *n*= 39. P values were calculated using the two-tailed *t*-test (**** represents p< 0.0001). Data are presented as mean ± SEM from three independent experiments. (E) RT-qPCR quantification of GPS2 target genes *AKAP1*, *ASNS*, *LONP1*, and *NDUFV1* in HeLa cells. Top row: si*CTL* and si*BAG6*, DMSO versus FCCP. Bottom row: si*CTL* and si*SQSTM1*, DMSO versus FCCP. Data normalized to β*-actin* and expressed relative to si*CTL* DMSO. P values were calculated using the two-tailed *t*-test (*represents p< 0.05; ** p< 0.01). Data are presented as mean ± SEM from three independent experiments. (F) Representative immunofluorescence images of HeLa cells transfected with control or KIF5B (si*KIF5B*) siRNA, stained for ATP5A (green), GPS2 (red), and DAPI (blue); single-channel GPS2 (gray scale) shown. The scale bar, 20 µm. Middle, quantification of nuclear GPS2 intensity. Right, representative immunoblot confirming KIF5B depletion, with β-tubulin as the loading control. Data points in the adjacent graph represent individual nuclei. For si*CTL*, *n*= 37; si*KIF5B*, *n*= 59. P values were calculated using the two-tailed *t*-test (**** represents p< 0.0001). Data are presented as mean ± SEM from three independent experiments. (G) RT-qPCR quantification of *AKAP1*, *ASNS*, *NDUFV1*, and *LONP1* in si*CTL* versus si*KIF5B* HeLa cells (no exogenous stressor). Data normalized to β*-actin* and expressed relative to si*CTL*. P values were calculated using the two-tailed *t*-test (*represents p< 0.05; ** p< 0.01). Data are presented as mean ± SEM from three independent experiments. (H) Representative immunofluorescence images of U2OS cells transfected with control or si*KIF5B* siRNA, stained for TOMM70 (green), GPS2 (magenta), and DAPI (blue); single-channel GPS2 (grayscale) shown. The scale bar represents 20 µm. Right, quantification of nuclear GPS2 intensity in U2OS cells. Data points in the adjacent graph represent individual nuclei. For si*CTL*, *n*= 16; si*KIF5B*, *n*= 19. P values were calculated using the two-tailed *t*-test (**** represents p< 0.0001). Data are presented as mean ± SEM from three independent experiments.

Having demonstrated that perinuclear clustering is necessary for efficient GPS2 retrograde signaling and that stress-induced perinuclear clustering is associated with MNCS formation, we asked whether modulating clustering affects MNCS formation. Indeed, we observed a significant decrease in the FCCP-induced increase in MNCS, as measured by SPLICS in U2OS cells, when mitochondria were prevented from clustering around the nucleus via BAG6 downregulation (**Figure 3A**). Conversely, KIF5B downregulation promotes a significant increase in the amount of MNCS observed in the absence of any exogenous stressor (**Figure 3B**).

**Figure 3.**
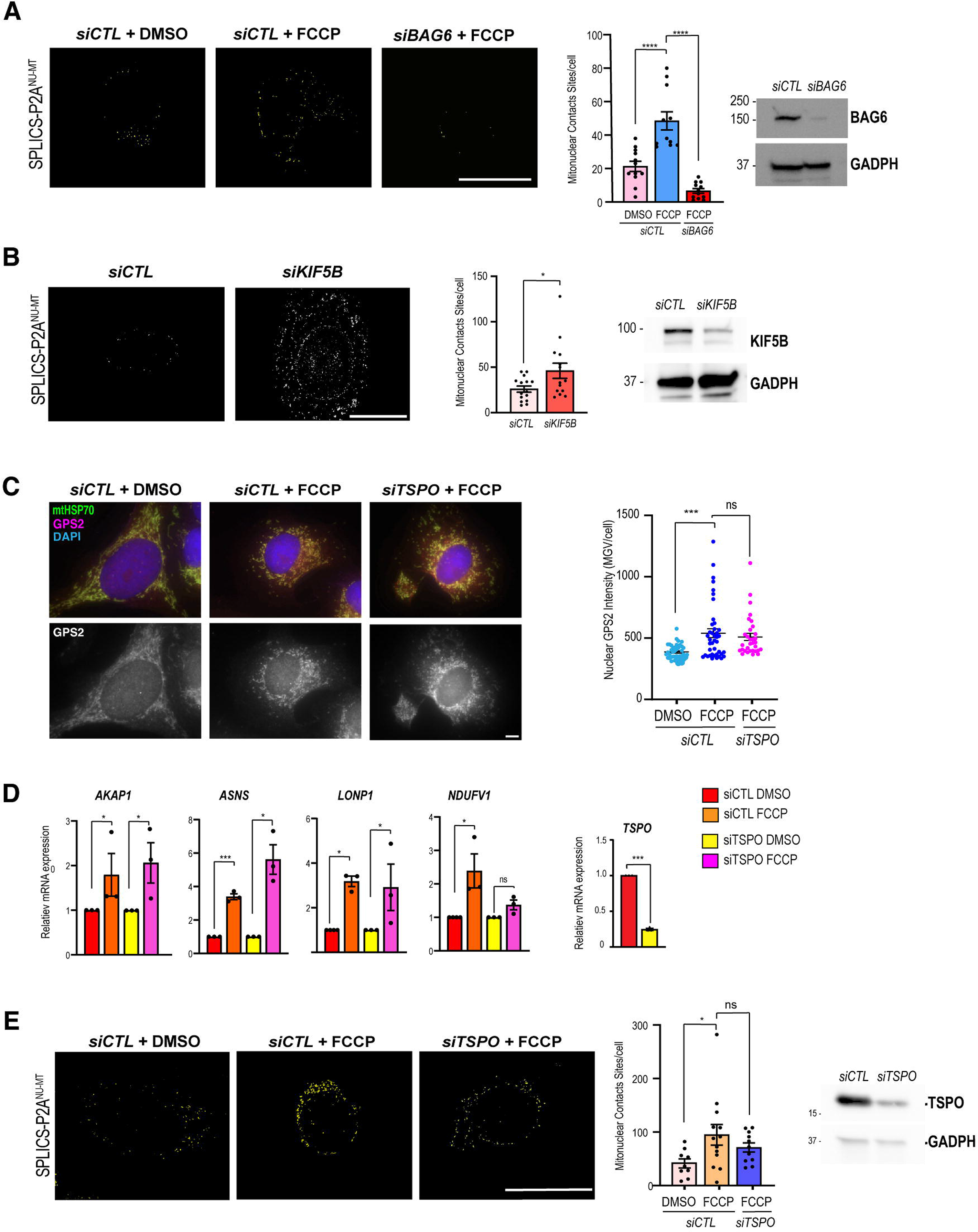
Perinuclear mitochondrial positioning regulates MNCS, whereas TSPO is dispensable for stress-induced contacts and GPS2 signaling. (A) MNCS quantified by the SPLICS-P2A^NU-MT^ reporter in U2OS cells: si*CTL*, si*CTL* + FCCP, and si*BAG6* + FCCP (3D-rendered reconstituted GFP, yellow). The scale bar, 20 µm. Right, quantification of MNCS per cell. Each dot represents one analyzed cell, identified by its corresponding nuclear ROI. Only cells with adequate SPLICS reporter expression and image quality for reliable three-dimensional rendering and contact-site quantification were included. For si*CTL*+DMSO, *n*= 12; si*CTL*+FCCP, *n*=11; si*BAG6* + FCCP, *n*=12. P values were calculated using the two-tailed *t*-test (* represents p< 0.05 and ** p< 0.01 respectively). Data are presented as mean ± SEM from three independent experiments. (B) MNCS were similarly quantified in si*CTL* and si*KIF5B* U2OS cells in the absence of an exogenous stressor. The scale bar, 20 µm. Right, quantification of MNCS per cell using the corresponding nuclear ROI. Each dot represents one analyzed cell. For si*CTL*, *n*= 15; si*KIF5B*, *n*=15. P values were calculated using the two-tailed *t*-test (* represents p< 0.05). Data are presented as mean ± SEM from three independent experiments. In (A) and (B), adjacent immunoblots confirm target depletion; GAPDH served as the loading control. (C) Representative immunofluorescence images of U2OS cells (si*CTL*+DMSO, si*CTL* + FCCP, si*TSPO* + FCCP) stained for mtHSP70 (green), GPS2 (magenta), and DAPI (blue); single-channel GPS2 (gray scale) shown. The scale bar, 20 µm. Right, quantification of nuclear GPS2 intensity (MGV per nucleus). Data points in the adjacent graph represent individual nuclei. For si*CTL*+DMSO, *n*= 41; si*CTL* + FCCP, *n*=41; si*TSPO* + FCCP, *n*=33. P values were calculated using the two-tailed *t*-test (*** represents p< 0.001; ns stands for non-significant). Data are presented as mean ± SEM from three independent experiments. (D) RT-qPCR quantification of *AKAP1*, *ASNS*, *LONP1*, and *NDUFV1* in si*CTL* versus si*TSPO* cells (DMSO versus FCCP), with *TSPO* knockdown validation (inset). Data normalized to β*-actin* and expressed relative to si*CTL* DMSO. P values were calculated using the two-tailed *t*-test (*** represents p< 0.001; ns stands for non-significant). Data are presented as mean ± SEM from three independent experiments. (E) MNCS quantified by SPLICS-P2A^NU-MT^ in U2OS cells: si*CTL* + DMSO, si*CTL* + FCCP, and si*TSPO* + FCCP. The scale bar, 20 µm. Right, quantification of MNCS per cell. Each dot represents one analyzed cell, identified by its corresponding nuclear ROI. Data points in the adjacent graph represent individual nuclei. For si*CTL* +DMSO, *n*= 9; si*CTL* + FCCP, *n*=13; si*TSPO* + FCCP, *n*=11. P values were calculated using the two-tailed *t*-test (* represents p< 0.05; ns stands for non-significant). Data are presented as mean ± SEM from three independent experiments.

Next, we asked whether MNCS are functionally required for retrograde signaling. Previous work revealed that TSPO, a translocator protein enriched at the outer mitochondrial membrane (OMM), acts as a molecular tether at the mitochondria-nucleus interface^42^, thus representing a prime candidate for modulating MNCS formation. Unexpectedly, we found that siRNA-mediated knockdown of TSPO did not impair GPS2 nuclear accumulation in response to FCCP (**Figure 3C**) nor the induction of GPS2 target genes, as assessed by RTqPCR (**Figure 3D**). While this indicates that TSPO is dispensable for GPS2 retrograde signaling, confirmation that TSPO downregulation impairs MNCS formation in our experimental model was required to conclude that MNCS are also dispensable. To this end, we explored the role of TSPO in MNCS formation in response to mitochondrial stress using the SPLICS-P2A^NU-MT^ reporter. As expected, TSPO knockdown resulted in a partial but significant reduction in MNCS; however, FCCP-induced contact sites remained detectable above the unstressed baseline (**Figure 3E**), in contrast to the near-complete loss observed upon downregulation of BAG6 (**Figure 3A**). These results indicate that while TSPO may contribute to a subset of contacts, it is not the primary mitochondrial molecular bridge for stress-induced MNCS in U2OS cells. We therefore reasoned that a different mitochondrial tether could be required for enabling GPS2 retrograde signaling via the formation of stress-induced MNCS.

Based on previous studies in yeast, in which Tom70-Cnm1 were identified as the first pair of MNCS tethers ^41^, and TOMM70 involvement in mitochondria-ER contact sites ^54^, we asked whether TOMM70 was required for mediating GPS2 retrograde signaling via MNCS. Indeed, knockdown of TOMM70 significantly impaired GPS2 nuclear translocation in response to FCCP (**Figure 4A**) and blunted the induction of GPS2 target genes (**Figure 4B**). Moreover, overexpression of TOMM70 led to a robust increase in MNCS, as quantified by the SPLICS-P2A^MT-NU^ reporter, even in the absence of any stress signal (**Figure 4C**), consistent with the properties expected for a tether ^24^. Together, these results indicate that TOMM70’s role as a yeast mitochondrial tether extends to mammalian cells and reveal that TOMM70 is required for stress-induced MNCS formation and functional coupling of MNCS and GPS2 retrograde signaling.

**Figure 4.**
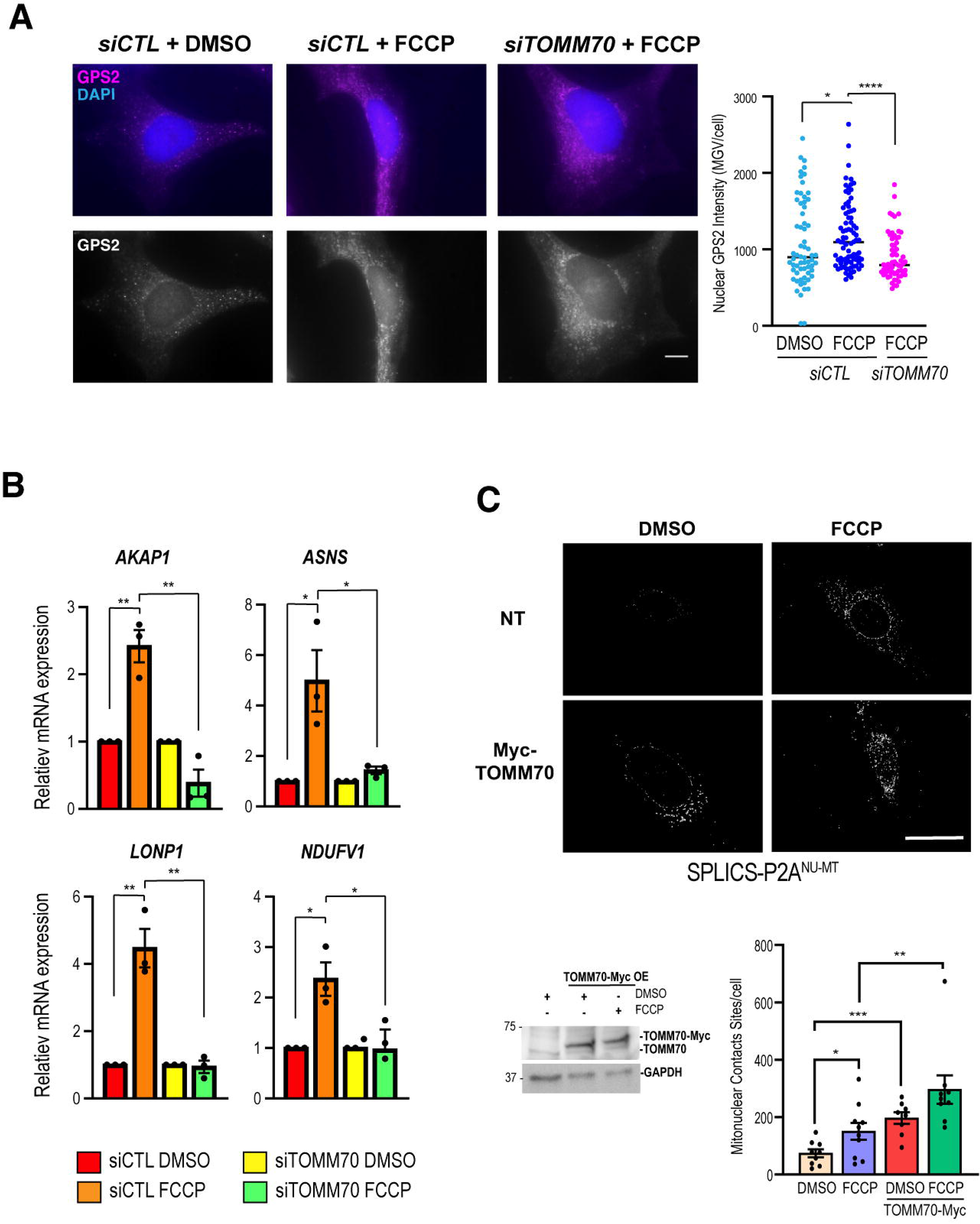
TOMM70 is required for stress-induced MNCS formation and GPS2 retrograde signaling. (A) Control and *TOMM70*-depleted HeLa cells were treated with DMSO or 10 µM FCCP for 1 h. si*CTL* + FCCP was taken as positive control. Representative immunofluorescence images stained for GPS2 (magenta), and DAPI (blue); single-channel GPS2 (gray scale) shown. The scale bar represents 20 µm. Right, quantification of nuclear GPS2 intensity in HeLa cells (MGV per nucleus). Data points in the adjacent graph represent individual nuclei. For si*CTL* + DMSO, *n*= 65; si*CTL* + FCCP, *n*= 82 and si*TOMM70* + FCCP, *n*= 64. P values were calculated using the two-tailed *t*-test (*represents p< 0.05 and **** p< 0.0001 respectively). Data are presented as mean ± SEM from three independent experiments. (B) RT-qPCR quantification of *AKAP1*, *ASNS*, *LONP1*, and *NDUFV1* in si*CTL* versus si*TOMM70* HeLa cells (DMSO versus FCCP). Data normalized to β*-actin* and expressed relative to si*CTL* DMSO. P values were calculated using the two-tailed *t*-test (* represents p< 0.05; ** p< 0.01). Data are presented as mean ± SEM from three independent experiments. (C) MNCS were quantified using SPLICS-P2A^NU-MT^ in control and TOMM70-Myc-overexpressing cells treated with DMSO or FCCP. Right, quantification of MNCS and immunoblot validation of TOMM70-Myc overexpression. The scale bar represents 20 µm. Right, quantification of MNCS per cell. Each dot represents one analyzed cell, identified by its corresponding nuclear ROI. Only cells with adequate SPLICS reporter expression and image quality for reliable three-dimensional rendering and contact-site quantification were included. For DMSO, *n*= 9; FCCP, *n*=10; TOMM70-Myc + DMSO, *n*=8 and TOMM70-Myc + FCCP, *n*=9. P values were calculated using the two-tailed *t*-test (*** represents p< 0.001; ** p< 0.01 and * p< 0.05 respectively). Data are presented as mean ± SEM from three independent experiments.

Having identified TOMM70 as a mitochondrial component of the MNCS used to mediate GPS2 retrograde signaling, we turned to the nucleus to characterize TOMM70’s complementary anchor. In yeast, the inner nuclear membrane protein Cnm1 mediates mitochondrial-nuclear tethering in association with Tomm70; however, no clear mammalian ortholog of Cnm1 has been identified, leading us to hypothesize that TOMM70 may interact with alternative partner(s). We reasoned that such a partner would likely interact with both TOMM70 and the retrograde cargo to be transported, and thus profiled GPS2-interacting proteins by IP/MS across fractionated cytosolic, mitochondrial, and nuclear protein extracts from WT and GPS2 KO mouse embryonic stem cells (MEFs^55^). To integrate the GPS2 interactomes with TOMM70-interacting proteins, we took advantage of published proximity proteomics studies performed in cells exposed to oxidative stress using the TOMM complex as bait. This revealed a strong enrichment of nuclear pore complex (NPC) components among both GPS2 and TOMM70/TOMM20 interactomes^56^. Notably, across GPS2 datasets we observed some noticeable differences. Mitochondrial, cytosolic, and nuclear datasets all included multiple components of the central NPC scaffold (NUP93, NUP155, NUP205). Mitochondrial and cytosolic datasets also included components of the outer NPC Y-ring (NUP107, NUP133) and RanBP2/NUP358. All these subunits were also identified as part of stress-induced TOMM20/TOMM70 proximity labeled datasets^56^. This was intriguing, as the juxtaposition of mitochondria and nuclear pores was first reported decades ago^34^, and recent studies have reported TOM and NPC coming together as a mitonuclear tether in Toxoplasma Gondii^57^.

The ultrastructural apposition of mitochondrial outer membranes and nuclear pores after a brief exposure to FCCP was confirmed in HeLa cells by transmission electron microscopy (**Figure 5A**). To investigate whether the NPC was required for retrograde signaling, we first downregulated the structural nucleoporin NUP93 and found a significant reduction in MNCS (**Figure 5B**), which supported the hypothesis that the NPC is required for the formation of mitonuclear contacts. The induction of GPS2 target genes was also significantly blunted upon NUP93 depletion (**Figure 5C**), indicating the NPC is required for GPS2-mediated retrograde signaling. Whether GPS2 signaling required the formation of MNCS, however, was not yet settled, as the loss of signaling upon NUP93 downregulation could simply reflect a role for the NPC in active transport rather than its requirement for structural tethering. Indeed, GPS2 harbors a nuclear localization signal, and its molecular weight places it near the threshold for NPC-dependent import. Treatment of HeLa cells with wheat germ agglutinin (WGA), which blocks active transport through the pore without disrupting its architecture, recapitulates the loss of GPS2 target gene activation observed with NUP93 knockdown (**Figure 5D**) due to the lack of stress-induced GPS2 nuclear accumulation (**Figure 5E**), further confirming that the NPC is required for GPS2 nuclear import. To investigate whether the NPC also serves a structural role in facilitating MNCS formation, we looked for NPC subunits that could selectively contribute to MNCS tethering. Based on its identification across TOMM20/TOMM70 and GPS2 mitochondrial interactomes, and its contribution to cytosolic filaments extending out into the cytosol, we focused on nucleoporin NUP358, also known as RanBP2. Indeed, we observed that downregulation of RanBP2 impaired MNCS formation, as quantified by SPLICS-PL2A^NU-MT^ (**Figure 6A**). Moreover, both GPS2 nuclear accumulation (**Figure 6B)** and target gene induction (**Figure 6C)** in response to FCCP were significantly impaired in the absence of RanBP2-mediated MNCS. Because the core scaffold of the NPC is not altered or made dysfunctional by RanBP2 depletion, and the general import/export of cargo across the pore remains functional^58–60^, which was recently confirmed by a concurrent study showing that RanBP2 downregulation prevents the formation of MNCS and nuclear transfer of ATP and phosphocreatine in cardiomyocytes without altering nucleus-cytosolic shuttling of protein cargo^45^, the failure in GPS2 nuclear import/signaling is interpreted as confirmation that RanBP2-based MNCS are essential for effective stress-induced retrograde signaling.

**Figure 5.**
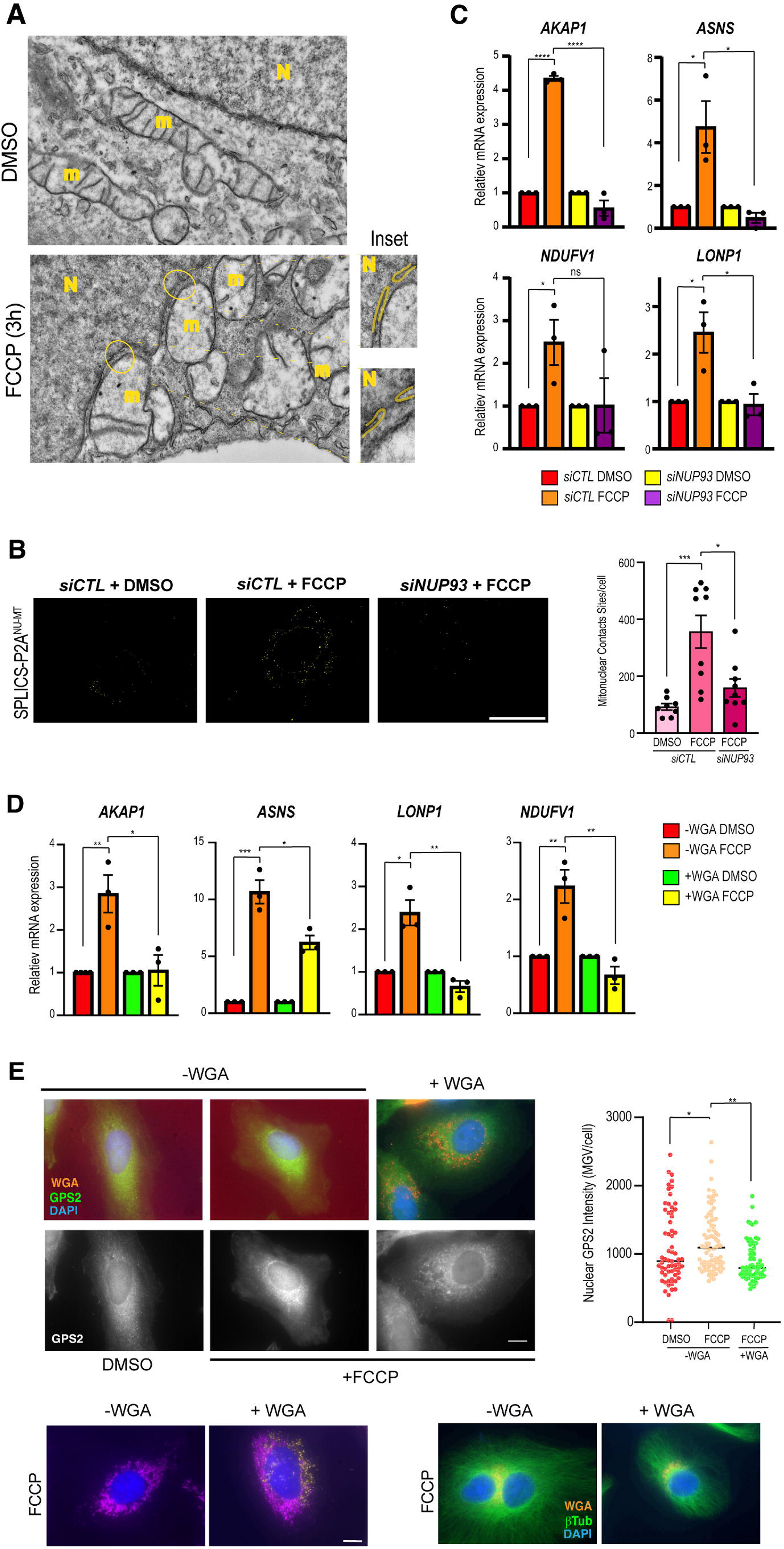
The nuclear pore complex is required for GPS2 nuclear import during retrograde signaling. (A) Representative transmission electron microscopy (TEM) images of HeLa cells treated with DMSO or FCCP (3 hours). N, nucleus; m, mitochondria. Insets highlight close apposition of mitochondrial outer membranes and nuclear pores at the nuclear envelope. Scale bars, 500 nm. (B) MNCS quantified by SPLICS-P2A^NU-MT^ in U2OS cells; reconstituted GFP signals are shown as 3D renderings: si*CTL* + DMSO, si*CTL* + FCCP, and si*NUP93* + FCCP. The scale bar represents 20 µm. Right, quantification of MNCS per cell. Each dot represents one analyzed cell, identified by its corresponding nuclear ROI. Only cells with adequate SPLICS reporter expression and image quality for reliable three-dimensional rendering and contact-site quantification were included. For si*CTL*+DMSO, *n*= 8; si*CTL* + FCCP, *n*=9; si*NUP93* + FCCP, *n*=9. P values were calculated using the two-tailed *t*-test (* represents p< 0.05 and ** p< 0.01). Data are presented as mean ± SEM from three independent experiments. (C) RT-qPCR analysis of the indicated GPS2 target genes in control and NUP93-depleted HeLa cells treated with DMSO or FCCP. All indicated genes were quantified from si*CTL* versus si*NUP93* HeLa cells (DMSO versus FCCP). Data normalized to β*-actin* and expressed relative to si*CTL* DMSO. P values were calculated using the two-tailed *t*-test (* represents p< 0.05; ** p< 0.01). Data are presented as mean ± SEM from three independent experiments. (D) RT-qPCR quantification of *AKAP1*, *ASNS*, *LONP1*, and *NDUFV1* in –WGA versus +WGA cells (DMSO versus FCCP). Data normalized to β*-actin* and expressed relative to DMSO with or without WGA. P values were calculated using the two-tailed *t*-test (* represents p< 0.05; ** p< 0.01). Data are presented as mean ± SEM from three independent experiments. (E) Representative immunofluorescence images of HeLa cells treated with DMSO or FCCP, without (–WGA) or with (+WGA) wheat germ agglutinin along with FCCP, stained for WGA (orange), GPS2 (green), and DAPI (blue); single-channel GPS2 (gray scale) shown. Bottom left, representative FCCP-treated cells with or without WGA used to assess perinuclear mitochondrial distribution. Bottom right, WGA, β-tubulin, and DAPI staining used to assess gross microtubule organization. (left) GPS2 (magenta)/DAPI ±WGA, and (right) WGA (orange)/β-tubulin (green)/DAPI ±WGA. Right, quantification of nuclear GPS2 intensity (MGV per nucleus). Data points in the adjacent graph represent individual nuclei. For -WGA + DMSO, *n*= 37; -WGA + FCCP, *n*= 39 and +WGA + FCCP, *n*= 40. P values were calculated using the two-tailed *t*-test (*represents p< 0.05 and ** p< 0.01 respectively). Data are presented as mean ± SEM from three independent experiments.

**Figure 6.**
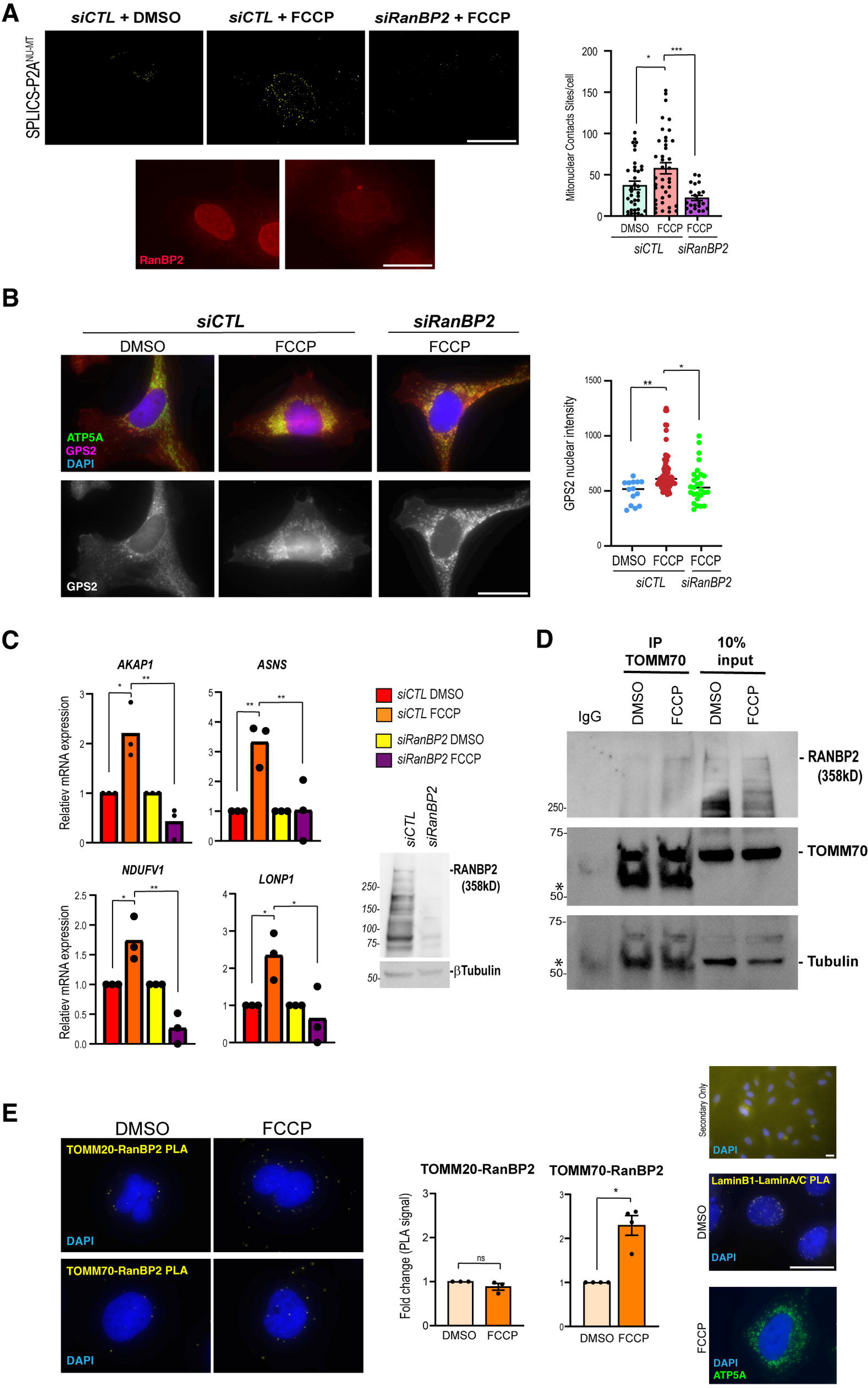
RanBP2/NUP358 is the nuclear anchor of MNCS and forms a stress-induced tether with TOMM70 that couples GPS2 to the nuclear pore. (A) MNCS quantified by SPLICS-P2A^NU-MT^ following FCCP treatment in control versus RanBP2-depleted U2OS cells. si*CTL* + DMSO, si*CTL* + FCCP, and si*RanBP2* + FCCP are the indicated experimental groups. Bottom, representative RanBP2 immunofluorescence confirming knockdown; RanBP2 is shown in red. The scale bar represents 20 µm. Right, quantification of MNCS per cell. Each dot represents one analyzed cell, identified by its corresponding nuclear ROI. Only cells with adequate SPLICS reporter expression and image quality for reliable three-dimensional rendering and contact-site quantification were included. For si*CTL*+DMSO, *n*= 36; si*CTL* + FCCP, *n*=40; si*RanBP2* + FCCP, *n*=24. P values were calculated using the two-tailed *t*-test (* represents p< 0.05 and *** p< 0.001). Data are presented as mean ± SEM from three independent experiments. (B) HeLa cells were treated with 10 µM FCCP for 1 hour, in control versus si*RanBP2* conditions. Si*CTL* + FCCP was taken as positive control. Representative immunofluorescence images stained for ATP5A (green), GPS2 (red), and DAPI (blue); single-channel GPS2 (gray scale) shown. The scale bar represents 20 µm. Right, quantification of nuclear GPS2 intensity in HeLa cells (MGV per nucleus). Data points in the adjacent graph represent individual nuclei. For si*CTL* + DMSO, *n*= 16; si*CTL* + FCCP, *n*= 66 and si*RanBP2* + FCCP, *n*= 43. P values were calculated using the two-tailed *t*-test (**represents p< 0.01 and *** p< 0.001 respectively). Data are presented as mean ± SEM from three independent experiments. (C) RT-qPCR quantification of GPS2 target genes (*AKAP1*, *ASNS*, *LONP1*, *NDUFV1*) in control versus si*RanBP2* cells (DMSO versus FCCP). Data normalized to β*-actin* and expressed relative to si*CTL* DMSO and si*RanBP2* DMSO respectively. P values were calculated using the two-tailed *t*-test (* represents p< 0.05; ** p< 0.01). Data are presented as mean ± SEM from three independent experiments. The adjacent blot represents the RanBP2 knock-down level. Here, β-tubulin is the loading control. (D) Reciprocal co-immunoprecipitation followed by immunoblotting demonstrates an association between TOMM70 and RanBP2. IgG served as immunoprecipitation control. Asterisks denote immunoglobulin heavy and light chains. β-tubulin served as the loading control for input lysates. (E) Representative images show the PLA for RanBP2–TOMM70 and RanBP2–TOMM20 pairs ± FCCP, showing the stress-induced increase is restricted to the RanBP2–TOMM70. Here, PLA puncta are shown in yellow and nuclei in blue (DAPI), Left. The quantification of PLA signals from both RanBP2–TOMM70 and RanBP2–TOMM20 pairs ± FCCP. The y-axis represents the fold change in PLA signal relative to the corresponding DMSO control. Lamin B1–lamin A/C PLA represents positive control; A primary antibody-only condition served as the negative control. ATP5A staining was used to visualize perinuclear mitochondrial distribution (Right). The scale bar represents 20 µm. Here, * stands for p<0.05 and ns represents non-significant. Data are presented as mean ± SEM from three independent experiments.

To further confirm the interaction between RanBP2 and TOMM70 at the mitonuclear interface, we performed pull-downs and proximity ligation assays (PLA). Immunoprecipitation of TOMM70 followed by western blot for RanBP2 confirmed a physical interaction between TOMM70 and RanBP2 that increases upon FCCP treatment (**Figure 6D**), thus providing biochemical evidence that these two proteins form a matched tether pair that bridges the mitochondrial outer membrane to the NPC scaffold in response to mitochondrial stress. For *in situ* visualization of the interaction between RanBP2 and the TOMM complex, we performed PLA with either TOMM70 or TOMM20. Interaction between the two sets of proteins was visualized and quantified as PLA dots by detecting the amplified circularized DNA with a fluorescent probe (DuoLink Technology). Unexpectedly, although both TOMM20 and TOMM70 reside within the TOM complex and exhibited basal proximity to RanBP2, we found that only the RanBP2– TOMM70 interaction increased significantly following mitochondrial stress (**Figure 6E**). Together, these results suggest that TOMM70’s role in the formation of MNCS is independent of its role as part of the import channel and reveal the TOMM70-RanBP2 tethering pair as a stress-responsive MNCS distinct from other mitonuclear contacts.

## Discussion

Our study resolves a long-standing question about perinuclear mitochondrial clustering: rather than a passive byproduct of mitochondrial dysfunction, the repositioning of stressed mitochondria toward the nucleus emerges as an active, required step in retrograde signaling. By using GPS2 nuclear translocation as a quantitative readout, we show that stress-induced perinuclear clustering, the formation of mitochondria-nucleus contact sites (MNCS), and the transcriptional activation of nuclear stress-response genes are mechanistically linked. Altering clustering in either direction, either by impairing it with p62/SQSTM1 or BAG6 depletion or by promoting it through KIF5B depletion, consistently alters GPS2 signaling output, highlighting the importance of spatial proximity as a controllable factor in the stress response. We also clarify the molecular structure of this interface: the outer-membrane receptor TOMM70 and the nucleoporin RanBP2/NUP358 serve as a tethering pair that assembles stress-responsive MNCS and is essential for delivering GPS2 to the nucleus. Conversely, the previously suggested tether TSPO is dispensable for GPS2 signaling. These findings collectively establish MNCS as an effective link connecting mitochondrial network structure with nuclear gene activity.

An important implication of our findings is that the nuclear pore complex is not merely a passive gateway for nucleocytoplasmic shuttling of cargo but an active participant in stress-induced mitonuclear communication. Supporting this model, depletion of the scaffold nucleoporin NUP93 markedly reduced MNCS, indicating that an intact nuclear pore architecture is required for the establishment or maintenance of these interfaces. Together with the requirement for RanBP2 in stress-induced retrograde signaling, these observations suggest that the nuclear pore complex provides a structural platform that spatially organizes mitonuclear communication. Within the NPC, RanBP2 is well positioned to play a central role due to its location extending out from the cytoplasmic face of the nuclear pore and its established roles as a multifunctional hub. Beyond its canonical role in nucleocytoplasmic transport, RanBP2 contains kinesin-and dynein-binding domains and has been shown to restrict ER–mitochondria connectivity through mTORC2/Akt ^61,62^. The convergence of motor-binding and membrane-tethering activities on a single scaffold is appealing because it enables a single molecule to coordinate two processes that our results indicate are essential for retrograde signaling and activation of the mitochondrial stress response: the microtubule-based delivery of mitochondria to the perinuclear space and their docking at the nuclear surface. In this view, RanBP2 acts as an integrator that reads mitochondrial position and converts it into a docking decision. Importantly, the recent demonstration that deletion of the RanBP2 C-terminal domain impairs MNCS and nuclear energetics without affecting bulk nuclear transport (PKA nuclear translocation, mRNA export, and RanGAP SUMOylation all remain intact)^45^ supports our conclusion that RanBP2’s contact-site function is genetically and biochemically separable from its transport function. This separability is what makes it plausible for a transport channel to double as a signaling tether without compromising essential traffic at the pore.

The identification of RanBP2 as a mitonuclear tethering platform across different experimental settings raises the question of how this nucleoporin differs from other NPC subunits. One key difference, in addition to positioning at the cytosolic edge of the pore, is that RanBP2 is an E3 SUMO ligase and a SUMO-interacting protein. Our previous studies show that GPS2 retrograde translocation is gated by SUMO^14^, and that the deSUMOylating enzyme SENP1, which promotes GPS2 retrograde translocation, is recruited to the NPC^63^. SUMOylation of mtHSP70/HSPA9 was also recently reported to regulate its export from mitochondria to the nucleus^64^. And Ulp, the yeast homolog of SENP proteins, was identified as a putative regulator of Cnm1-Tomm70 contact sites in S. Cerevisiae^41^. Together, these findings suggest that SUMO cycling at MNCS is a key component of mitonuclear communication. Within this framework, the buildup of mitochondrial proteins on the cytosolic side of the OMM, which occurs when depolarization blocks the TOM channel, may be crucial for establishing their SUMO status and directing them to the nucleus. TOMM70 would therefore function as a sensor, and its binding to stalled, SUMO-modified cargo could trigger the RanBP2 contact, facilitating the transfer of that cargo to the pore, where its de-sumoylation by SENP1 is coupled to import. Notably, SENP1 activity is directly inhibited by L-lactate, which binds zinc in the enzyme’s active site^65^, and is coupled to nutrient sensing through AMPK^66^. Thus, placing SENP1 at the heart of the retrograde switch would directly link mitonuclear communication to the cell’s metabolic state, possibly helping dissect how mitochondrial stress responses and mitonuclear communication pathways are differently tuned across cell types and metabolic conditions.

The identification of TOMM70, rather than the core translocon, as a tethering partner for RanBP2 in mediating stress-induced contacts is consistent with a growing appreciation that TOMM70 has moonlighting functions independent of its role as an import receptor^54,67–69^. TOMM70 is a TPR-domain adaptor that docks cytosolic chaperones and independently supports ER-mitochondria calcium transfer via IP3R3, MAVS-dependent antiviral signaling, and PINK1/Parkin recruitment. In yeast, Tom70 was identified as the mitochondrial tether for Cnm1-mediated nucleus–mitochondria contacts^41^, and a recent study shows that Tom70 organizes mitochondria–nuclear-envelope contacts governing nuclear pore inheritance during gametogenesis in a manner explicitly independent of its import activity^70^. Our finding that TOMM70’s tethering role is separable from the general import channel, as the stress-induced RanBP2 interaction is TOMM70-specific and not shared by TOMM20, extends this import-independent function to mammalian retrograde signaling and suggests that ancestral Tom70– nuclear pore contacts have been repurposed to gate cargo, not merely to position the organelle.

During the preparation of this manuscript, an independent study identified RanBP2–VDAC1 contacts that support mitochondria–nucleus communication by facilitating nuclear energetics^45^. Our findings identify TOMM70 as a RanBP2 partner required for stress-induced retrograde signaling. Together, these findings suggest that RanBP2 can participate in mechanistically distinct mitonuclear interfaces with different biological functions: the constitutive VDAC1– RanBP2 contacts supply energy metabolites to sustain nuclear ATP-dependent processes, whereas stress-inducible TOMM70–RanBP2 contacts provide a distinct route for transporting protein cargo, such as GPS2, during the retrograde response. Notably, this interpretation aligns with an independent SPLICS-based dissection of MNCS in HeLa cells, which found no single dominant tether but rather a distributed, dynamic network in which TOMM70 is the strongest single contributor to contact abundance^40^. A tether repertoire specialized by cargo and cell status (energy versus protein, basal versus stress) would let the cell scale the interface to demand and may reconcile the apparently competing molecular identities reported for this contact.

A clear implication of using key transport machinery as a structural tether is that the contact cannot be permanent. Unlike other contacts, which can be long-lived, an MNCS built upon the NPC must be transient and reversible, as nuclear pores locked into a mitochondrial contact would be occluded for bulk traffic. We therefore anticipate that MNCS are dynamically regulated, forming rapidly upon stress and dissolving once the retrograde signal has been transmitted, and that their turnover kinetics, rather than their steady-state abundance, may be the physiologically relevant variable. Defining what triggers assembly and disassembly, and whether individual pores cycle between transport-competent and tether-engaged states, is an important question for future work, and may reveal that the biophysical nature of the mitonuclear contacts differs fundamentally from that of other inter-organelle junctions.

Finally, our perturbations reveal a functional role for spatial reorganization of the mitochondrial network in retrograde signaling, showing that mitochondrial clustering around the nucleus is essential for activation of stress response genes. In accordance with prior studies, our results confirm that stress-induced perinuclear collapse depends on p62/SQSTM1 and BAG6, whereas constitutive dispersal of the network toward the periphery depends on the kinesin KIF5B, whose loss is sufficient to drive mitochondria onto the nucleus and to activate GPS2 signaling even in the absence of an exogenous stressor. This raises the question of whether mitochondrial position could also gate mitonuclear communication in physiological, non-stress settings where network geometry is intrinsically polarized, such as in neurons, where perinuclear versus axonal mitochondrial pools are tightly controlled, or during differentiation. While the stress-specific p62/BAG6/KIF5B logic is unlikely to be relevant in physiological contexts, dissecting the role of canonical adaptors that regulate mitochondrial movement towards the plus-and minus-end of microtubules, such as Miro1/2 and TRAK1/2 ^71–73^, in the formation of MNCS may reveal an additional layer of spatial regulation of mitonuclear communication.

Overall, our results suggest that the position of mitochondria, their docking onto the nucleus, and the metabolic signals they transport are collectively integrated at the nuclear pore, influencing the transcriptional response to mitochondrial stress.

## Acknowledgements

We thank Dr. Jomon Joseph and Dr. Mary C. Dasso for generously providing the RanBP2 expression construct, Dr. Paola Pizzo for the TOMM70 expression construct, and Dr. Tito Calì for the SPLICS-P2A^NU-MT^ reporter. We acknowledge Dr. Andrew Emili and Dr. Julian Kwan at the Center for Network Systems Biology (CNSB) for their assistance with MS-based studies of the GPS2 interactome; and Dr. Maria Ericsson at the Harvard Electron Microscopy Core for processing of TEM images. We are grateful to Stefan Isaac and all members of the Isaac lab for insightful discussions and valuable scientific input throughout the course of this study. We also thank Dr. Bob Varelas and all lab members for providing access to essential laboratory equipment and technical resources. This work was supported by the National Institute of General Medical Sciences (NIGMS), National Institutes of Health, under award R35GM149339 to VP.

## Experimental Methods

### Cell culture & drug treatments

HeLa (CCL2), U2OS and 3T3-L1 cells were purchased from ATCC (Manassas, VA, USA). HeLa and U2OS cells were grown in complete DMEM (Thermo Scientific, Cat. No.: 11965118) with 10% Fetal Bovine Serum (FBS) (Gibco, Cat. No.: A5256801), while 3T3-L1 was maintained in complete DMEM with 10% bovine calf serum (Sigma Millipore, Cat. No.: 12133C). All cultures were grown with 10% Pen-strep (Corning, Cat. No.: 30-002-CI) except for 3T3-L1 transfection. Cells were treated for 1 hour or 3 hours with 10 µM FCCP, 1 µM Rotenone, 1 µM Oligomycin or 50 µM MitoBloCK-6.

### Plasmids

SPLICS-P2A^NU-MT^ (Addgene: 210129) was used for the mito-nuclear contact sites evaluation. SPLICS MT-ER Short P2A (Addgene: 164108) was used to identify endoplasmic reticulum-mitochondria (ER-MT) contact sites. Myc-TOMM70 mammalian expression vector was a generous gift from Dr. Paola Pizzo^54^.

### Plasmid and siRNA transfection

For SPLICS-P2A^NU-MT^ reporter transfection, cells were grown on coverslips in 12-well plates at 5*10^4^ cell density per well in complete DMEM. Reporter transfection was carried out using 200 ng of SPLICS-P2A^NU-MT^ and 1 µg of SPLICS-P2A^MT-ER^ with Lipofectamine 2000 (Thermo Fisher, Cat. No.: 11668027). All transfections were carried out in reduced serum medium (OptiMEM, Fisher Scientific, Cat. No.: 31985070) for 4 hours, after which the OptiMEM was removed, and the cells were fed with complete medium to allow growth for the next 36 hours. Transient siRNA transfections were carried out with Lipofectamine 3000 (Thermo Fisher, Cat. No.: L3000008) following the manufacturer’s instructions. Cells were seeded in either 6-well (for WB and RTqPCR) or 12-well (Immunofluorescence) plates at 50-60% confluency and transfected the following day with 20 pmol. Transfection was carried out, without changing the medium, for 24 hrs (si*NUP93* and si*Bag6*); 48 hrs (si*p62/SQSTM1* and si*TOMM70*); 72 hrs (si*TSPO*, si*RanBP2* and si*Kif5B*) to achieve maximum knockdown of the gene of interest.

### Immunofluorescence & Antibodies

Cells were grown on coverslips (Chem Glass Life Sciences, USA) and treated accordingly to the experimental setting of each experiment. Fixation was done with 4% paraformaldehyde diluted in 1X phosphate buffer solution (PBS) for 15 min at room temperature (RT) with shaking. Cells were permeabilized with 0.3% Triton-X for 5 min at RT, followed by a blocking step (0.5% BSA + 1:500 Donkey Serum in 1X PBS) for 1 h at RT. Cells were incubated with the primary antibodies (1:100, diluted in blocking solution) overnight at 4 degrees in a humid chamber. The following day, after three 10 min washes with 1X PBS on the shaker, cells were incubated with secondary antibodies (1:500, diluted in blocking solution) for 1 h at RT, followed by subsequent washes with 1X PBS. Cells were then counterstained with 1 mM DAPI (Molecular Probes, Oregon, USA) for 10 min at RT. In all immunofluorescence experiments, we included a control group (stained only with the secondary antibody) that was used for background fluorescence adjustment.

For IF, we used the following primary antibodies: mouse α-ATP5A (Invitrogen, 43-9800); mouse α-mtHsp70 monoclonal antibody JG1 (Invitrogen, MA3-028); mouse α-TOM70 (Santa Cruz, sc-390545); rabbit α-TOM70 (Proteintech, 14528-1-AP); rabbit α-TOM20 (Proteintech, 11802-1-AP); mouse α-RanBP2 (Santa Cruz; sc-74518); mouse α-NUP93 (Santa Cruz; sc-374400); mouse α-Bag6 (Santa Cruz; sc-365928); mouse α-p62 (EMD Millipore; MABC32); rabbit α-Kif5B (Abcam, AB167429); rabbit α-TSPO (Abcam, AB109497); mouse α-GAPDH (Santa Cruz; sc-47724); mouse α-beta tubulin (Sigma, D66); rabbit α-GPS2 (Ct) (Custom, aa1-20); rabbit α-Lamin A/C (Proteintech; 10298-1-AP). Secondary antibodies included goat α-mouse Alexa Fluor 488 (Invitrogen) and goat α-mouse Alexa Fluor 647 (Invitrogen).

### Proximity ligation assay

Proximity ligation assays (PLA) were performed using the Duolink In Situ Detection Orange Kit (mouse/rabbit; Sigma-Aldrich, Cat. No.: DUO92102-1KT) according to the manufacturer’s instructions. Samples were processed according to the IF procedure described above, including overnight incubation with primary antibodies. The following day, cells were rinsed with the kit-provided Buffer A and incubated with PLA probe-conjugated secondary antibodies for 1 h at 37°C. DNA probes in proximity were subsequently ligated by incubation with the supplied DNA ligase for 30 min at 37°C. Signal amplification was then performed by rolling-circle amplification using the supplied DNA polymerase for 100 min at 37°C. Following the prescribed washing steps with the kit-provided buffers, samples were mounted using the DAPI-containing mounting medium included in the kit, coverslips were sealed with nail polish and stored protected from light until imaging. Quantification was performed following published protocol^74^.

### Microscopy

Cells were visualized using an Olympus BX61 microscope equipped with a spinning disc confocal scanning unit. For SPLICS reporter imaging, images were acquired as three-dimensional optical stacks with a 0.3-µm z-step size (0.3 µm per optical slice) using an ORCA-ER charge-coupled device (CCD) camera (C4742-80; Hamamatsu Photonics, Hamamatsu, Japan) controlled by MetaMorph software (Molecular Devices, San Jose, CA, USA). For all other imaging experiments, single-plane images were acquired. All images were collected using either a 100× oil-immersion objective (NA 1.4) or a 20× objective. Coverslips were mounted with ProLong™ Gold Antifade Mountant (Thermo Fisher Scientific, Waltham, MA, USA). Images acquired with the 100× objective were obtained using Olympus immersion oil. All images were acquired at room temperature. Image processing and analysis were performed using ImageJ/Fiji. SPLICS images were analyzed as previously described^36^.

### Transmission electron microscopy (TEM)

Cells were grown on coverslips (Chem Glass Life Sciences, USA) and fixed in a solution of 0.2% EM-grade Glutaraldehyde (Electron Microscopy Sciences) and 4% EM-grade paraformaldehyde (Electron Microscopy Sciences) for 30 min. Fixed cells were washed with 0.1 M sodium cacodylate buffer and post-fixed with 1% osmium tetroxide/1.5% potassium ferrocyanide (in H2O) for 2 hrs. Samples were then washed in a maleate buffer and post-fixed in 1% uranyl acetate in maleate buffer for 1 hr. Samples were then rinsed in ddH20 and dehydrated through a series of ethanol (50%, 70%, 95%, (2x)100%) for 15 minutes per solution. Dehydrated samples were put in propylene oxide for 5 minutes before they were infiltrated in Epon mixed 1:1 with propylene oxide overnight at 4C. Samples were polymerized in a 60C oven in Epon resin for 48 hours. They were then sectioned into 80nm thin sections and imaged on a JEOL 1200EX Transmission Electron Microscope with an AMT 2k CCD camera.

### Quantitative reverse transcription PCR (RT-qPCR)

Approximately 500 ng of total RNA isolated from human cells was reverse transcribed in a 20 µL reaction using the iScript™ cDNA Synthesis Kit (Bio-Rad, Cat. No. 1708890) according to the manufacturer’s instructions. The resulting cDNA was diluted and used as template for quantitative PCR using PowerTrack™ SYBR Green Master Mix (Thermo Scientific, Cat. No. A46110) on an Applied Biosystems QuantStudio™ 7 Pro Real-Time PCR System (Thermo Fisher Scientific, Waltham, MA, USA). Cycle threshold (Ct) is determined automatically by the QuantStudio™ Design and Analysis Software based on the fluorescence amplification curve. Relative mRNA expression levels were calculated using the comparative Ct (ΔΔCt) method and normalized to the housekeeping gene β*-actin*. Graphs were generated using GraphPad Prism software.

The following primer pairs were used for RT-qPCR:

human *AKAP1* F: CCAAGTGGTTGCCTCCTACG

human *AKAP1* R: TTTGCCGGAGCACGTCTACT

human *ASNS* F: GATGTACCCCTGCACGCCCT

human *ASNS* R: AGGATCCTGAGGTTGTTC

human *NDUFV1* F: TGAGACGGTGCTGATGGACTTC

human *NDUFV1* R: AGGCGGGCGATGGCTTTC

human *Lonp1* F: CCTGACTGCAGAGATCGTGA

human *Lonp1* R: CCCATGTCGCTCAGGTAGAT

human *TSPO* F: TCTTTGGTGCCCGACAAAT

human *TSPO* R: GGTACCAGGCCACGGTAGT

human β*-actin* F: CCTCGCCTTTGCCGATCC

human β*-actin* R: GCGCGGCGATATCATCATCC

### Immunoblotting and Co-Immunoprecipitation

For whole cell extracts, cell pellets were lysed in ice-cold RIPA buffer (50 mM Tris-HCl, pH 7.5; 150 mM NaCl; 1% Triton X-100; 0.1% SDS) or IPH buffer (50 mM Tris-HCl, pH 8.0; 150 mM NaCl; 5 mM EDTA; 0.5% NP-40), in the presence of protease and phosphatase inhibitors. Lysates were incubated for 30 min in the cold room under gentle shaking to facilitate lysis, and then centrifuged at 13,500 rpm for 15 min using an Eppendorf 5424 R refrigerated benchtop microcentrifuge (Eppendorf AG, Hamburg, Germany), after which the supernatants were collected and protein concentration was determined using a Pierce™ BCA Protein Assay Kit (Thermo Fisher Scientific, Waltham, MA, USA) and quantified using a Thermo Scientific™ NanoDrop™ 2000c (Thermo Fisher Scientific, Waltham, MA, USA). Protein lysates were adjusted to equal concentrations and mixed with 1× SDS sample buffer (Bio-Rad Laboratories, Hercules, CA, USA), followed by denaturation at 98°C for 5 min. Equal amounts of protein were resolved on 10% polyacrylamide gels and transferred onto 0.22-µm pore-size Immobilon®-P polyvinylidene difluoride (PVDF) membranes (Millipore Sigma, Burlington, MA, USA) using either a Trans-Blot® Turbo™ Transfer System (Bio-Rad Laboratories, Hercules, CA, USA) according to the manufacturer’s instructions or by conventional wet transfer. For RanBP2 protein analysis, proteins were resolved on 4–15% polyacrylamide gradient gels, followed by overnight wet transfer. Membranes were then rinsed with PBS containing 0.1% Tween-20 (PBS-T), blocked in 3% nonfat dry milk prepared in PBS-T for 1 h at room temperature, and incubated overnight at 4°C with primary antibodies diluted in 0.1% PBS-T under gentle rotation. Following primary antibody incubation, membranes were washed with PBS-T and incubated with the corresponding horseradish peroxidase (HRP)-conjugated secondary antibodies (Cell Signaling Technology, Danvers, MA, USA). Immunoreactive bands were visualized using Bio-Rad Clarity™ Western ECL Substrate (Bio-Rad Laboratories, Hercules, CA, USA) and imaged with a ChemiDoc™ Imaging System (Bio-Rad Laboratories, Hercules, CA, USA). When reprobing was required, membranes were treated with a commercial stripping buffer (Thermo Fisher Scientific, Waltham, MA, USA) according to the manufacturer’s instructions before repeating the blocking and immunoblotting procedure.

## Co-immunoprecipitation

For IP, cells were lysed directly on the culture dish by the addition of IPH buffer, and protein concentrations were determined using a Pierce™ BCA Protein Assay Kit (Thermo Fisher Scientific, Waltham, MA, USA) according to the manufacturer’s instructions. 500 μg to 2 mg of total protein was incubated with the appropriate primary antibody overnight at 4°C under gentle rotation to immunoprecipitate the protein of interest. The following day, 30 μL of Protein A-or Protein G-coupled bead slurry (Invitrogen™, Thermo Fisher Scientific, Waltham, MA, USA), selected according to the host species and IgG subtype of the primary antibody, was added to each sample and incubated for 1 h at 4°C under gentle rotation. Following incubation and washes bound proteins were eluted by adding 2× SDS sample loading buffer (Bio-Rad Laboratories, Hercules, CA, USA) and boiling the samples at 95°C for 10 min. The eluted proteins were subsequently resolved by SDS-PAGE on polyacrylamide gels and analyzed by WB.

### WGA Microinjection

Hela cells were grown on coverslips (Chem Glass Life Sciences) suitable for microinjection until they reached a confluency of 50%. The following day, we microinjected the cells with 400μg/ml WGA (WGA FAlexa fluor 555; Thermo Scientific; W32464) for 6 hours using the 4D-Nucleofector method (Lonza 4D-Nucleofector™ System, Program number CN-114). After WGA administration, we treated cells with FCCP for 1 hour (imaging) & 3 hours (RT-qPCR), respectively. After treatment, cells were fixed and continued with the staining protocol described above. For RNA extraction, the above protocol was followed.

### Images Quantification

Mitochondria–nucleus contact sites were quantified using Fiji^75^ with the VolumeJ plugin and custom ImageJ macros for contact-site analysis, as described previously^36^. For each probe-positive cell, a region of interest encompassing the nucleus was manually defined so that only contact sites localized at the nuclear envelope were included in the analysis. All images were processed using identical analysis parameters, and contact site measurements were normalized to the corresponding control samples.

Intracellular distance measurements were also quantified using FIJI^75^. The minimum distance between each mitochondrion and the nuclear envelope was determined by drawing a straight line from the mitochondrial edge nearest the nucleus to the edge of the nuclear envelope and recording the line length in pixels. Pixel-to-micrometer (μm) conversion was performed by measuring, in pixels, the scale bar automatically generated by the image acquisition software and using this value to calibrate all recorded distances.

To quantify the mean gray value of GPS2 fluorescence, we used ImageJ. Measurements were performed within the nuclear region, using the nuclear focal plane while excluding any planes containing mitochondrial signal to ensure accurate quantification of nuclear GPS2 intensity. To calculate fold changes in nuclear GPS2 intensity, the mean gray value of each experimental condition was normalized to that of the corresponding control samples. Statistical significance was assessed using a two-tailed *t*-test unless otherwise indicated.

## Declaration of generative AI and AI-assisted technologies in the manuscript preparation process

During the preparation of this work, the authors used Grammarly and ClaudeCowork for polishing language and improving readability. The authors reviewed and edited the output as needed and take full responsibility for the content of the published article.

---

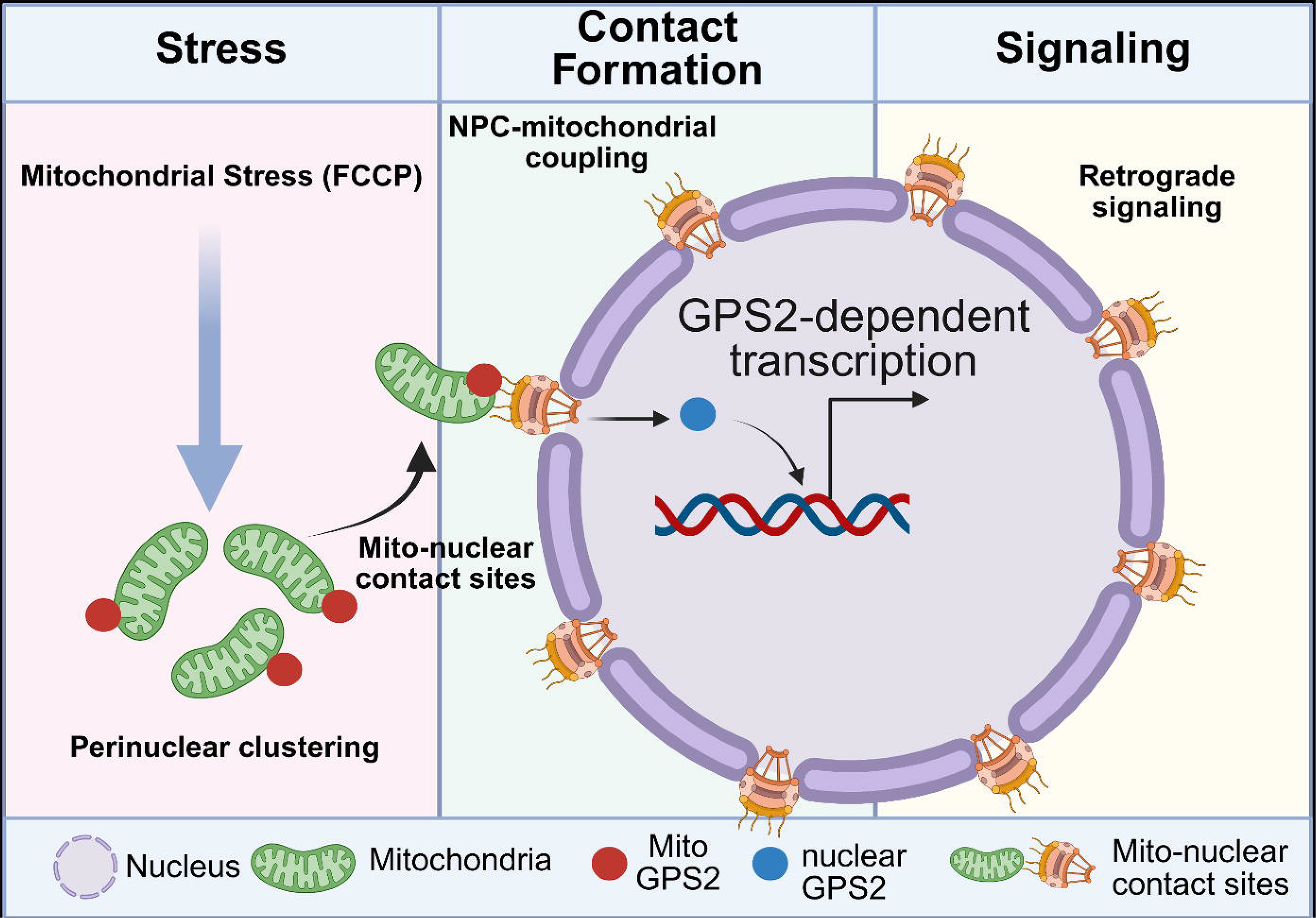

**Supplemental Figure S1.**
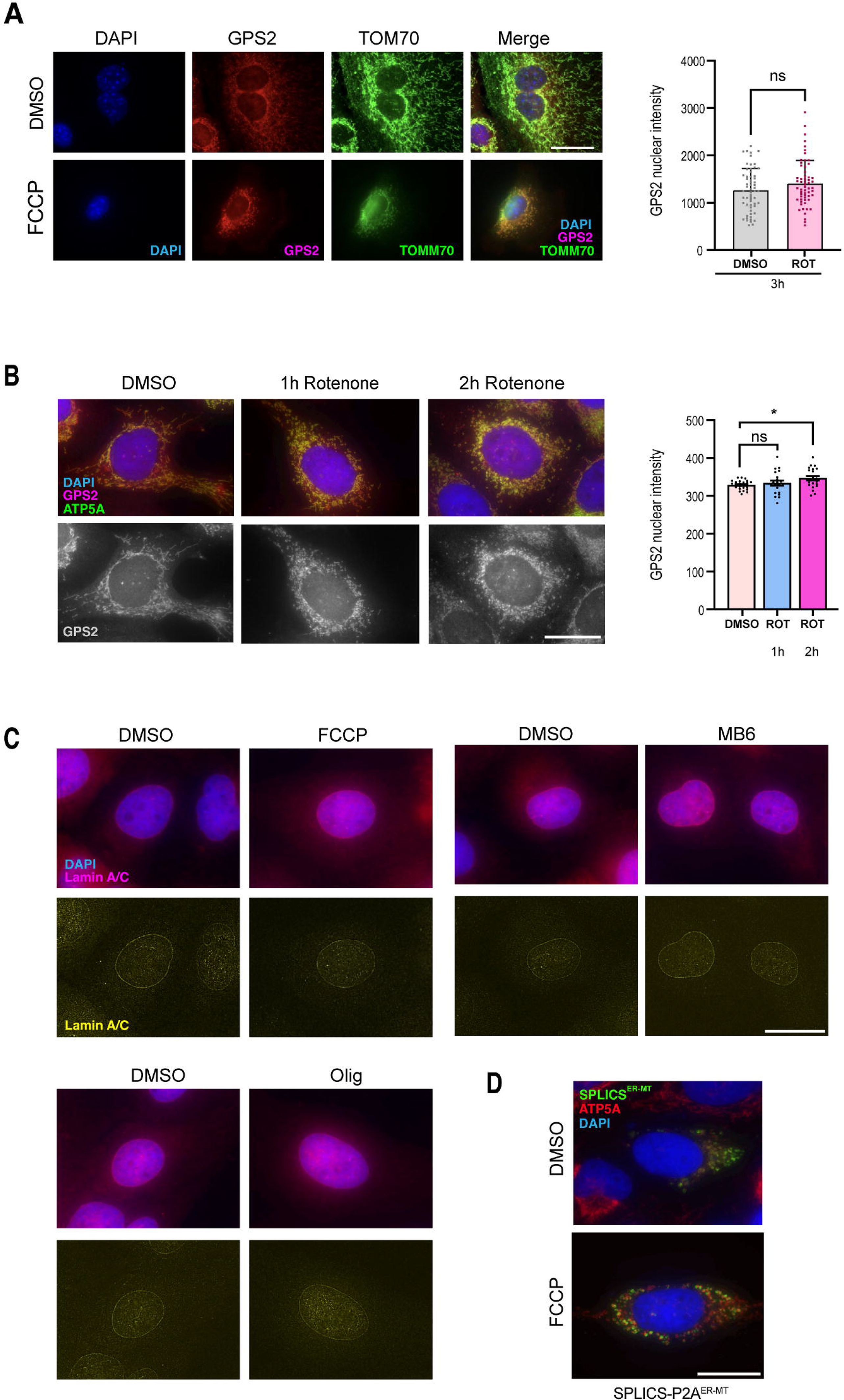
**Kinetics of GPS2 nuclear accumulation in response to Complex I inhibition by Rotenone treatment, nuclear envelope integrity, and mitochondria–ER contact sites. Related to Figure 1.** (A) Representative immunofluorescence images of 3T3-L1 cells treated for 3 hours with Rotenone 1µM or DMSO control, stained for DAPI (blue), GPS2 (red), and TOMM70 (green). Right, quantification of nuclear GPS2 intensity (MGV per nucleus), DMSO versus ROT (3 h). Statistical analysis of three independent experiments was performed using an unpaired t-test. Statistical significance was defined as follows: p-value 0.1234 (ns), 0.0332 (*), 0.0021 (**), 0.0002 (***), and <0.0001 (****). (B) Representative immunofluorescence images of U2OS cells treated with DMSO, 1 hour Rotenone, or 2 hours Rotenone, stained for DAPI (blue), GPS2 (magenta), and ATP5A (green); single-channel GPS2 (gray scale) shown. The scale bar represents 20 µm. Data points in the adjacent graph represent individual nuclei for the evaluation of mean nuclear GPS2 intensity. For DMSO, *n*= 25; 1 hour Rotenone, *n*=21; 2 hours Rotenone, *n*=22. P values were calculated using one-way ANOVA test (* represents p< 0.05 and ns stands for non-significant). Data are presented as mean ± SEM from three independent experiments. (C) Lamin immunofluorescence demonstrating that nuclear envelope structure is not grossly altered by mitochondrial stress. Top, merged lamin A/C and DAPI images. Bottom, identically processed lamin A/C images displayed to improve visualization; the same convolution and Gaussian-blur parameters were applied to all conditions. The scale bar represents 20 µm. (D) Short-range Mitochondria–ER contact sites were visualized by the SPLICS-P2A^ER-MT^ reporter following FCCP treatment, showing no stress-induced increase in perinuclear mito-ER contacts. Representative channels show SPLICS-GFP (green), DAPI (blue), and ATP5A (red). Scale bar: 20 µm.

